# C. elegans BRC-1-BRD-1 functions at an early step of DSB processing and inhibits supernumerary crossovers during male meiosis

**DOI:** 10.1101/2020.04.27.064097

**Authors:** Qianyan Li, Sara Hariri, JoAnne Engebrecht

**Affiliations:** Department of Molecular and Cellular Biology, and Biochemistry, Molecular, Cellular and Developmental Biology Graduate Group, University of California, Davis

**Keywords:** BRC-1-BRD-1, crossovers, meiosis, recombination, sex

## Abstract

Meiosis is regulated in a sex-specific manner to produce two distinct gametes, sperm and oocytes, for sexual reproduction. To determine how meiotic recombination is regulated in spermatogenesis, we analyzed the meiotic phenotypes of mutants in the tumor suppressor E3 ubiquitin ligase BRC-1-BRD-1 complex in *Caenorhabditis elegans* male meiosis. Unlike in mammals, this complex is not required for meiotic sex chromosome inactivation, the process whereby hemizygous sex chromosomes are transcriptionally silenced. Interestingly, *brc-1* and *brd-1* mutants showed meiotic recombination phenotypes that are largely opposing to those previously reported for female meiosis. Fewer meiotic recombination foci marked by the recombinase RAD-51 were observed in *brc-1* and *brd-1* mutants, and the reduction in RAD-51 foci can be suppressed by mutation of nonhomologous end joining proteins. We show that concentration of BRC-1-BRD-1 to sites of meiotic recombination is dependent on DNA end resection, suggesting that BRC-1-BRD-1 regulates the processing of meiotic double strand breaks to promote repair by homologous recombination, similar to a role for the complex in somatic cells. We also show that BRC-1-BRD-1 is important to promote progeny viability when male meiosis is perturbed by mutations that block the pairing and synapsis of different chromosome pairs, although the complex is not required to stabilize the RAD-51 filament as in female meiosis under the same conditions. Analyses of crossover designation and formation reveal that BRC-1-BRD-1 inhibits supernumerary crossovers when meiosis is perturbed. Together, our findings suggest that BRC-1-BRD-1 regulates different aspects of meiotic recombination in male and female meiosis.

## Introduction

Meiosis is essential for sexual reproduction and results in the precise halving of the genome for packaging into gametes. Chromosomes must be accurately segregated during meiosis to ensure that the next generation has the correct genomic complement. In metazoans with defined sexes, the products of meiosis, sperm and oocytes, contribute not only haploid genomes but also unique cellular components to support embryonic development. In addition to the striking morphological differences between sperm and oocytes, the process of meiosis itself exhibits extensive sexual dimorphism with respect to the temporal program of events, the extent and placement of recombination, checkpoint signaling, chromosome segregation and sex chromosome behavior (Morelli and Cohen 2005; Turner 2007; Nagaoka *et al.* 2012; Bury *et al.* 2016; Cahoon and Libuda 2019). However, the underlying mechanisms governing these differences are not well understood.

Meiotic chromosome segregation relies on establishing connections between homologous chromosomes. In most organisms this is accomplished by the intentional induction of hundreds of double-strand breaks (DSBs) by the conserved topoisomerase Spo11 and accessory proteins (Keeney *et al.* 1997; Dernburg *et al.* 1998). A subset of meiotic DSBs use a non-sister chromatid as template for repair by homologous recombination (HR) to generate crossovers that ensure disjunction and promote genetic variation. In almost all animals and plants where it has been examined, crossovers differ in number, placement and spacing in the sexes (Lenormand and Dutheil 2005; Gruhn *et al.* 2013; Stapley *et al.* 2017; Kianian *et al.* 2018; Lloyd and Jenczewski 2019).

Knowledge is lacking with respect to the contributions of different pathways to repair of DSBs not destined to form crossovers and whether their use differs in the sexes. During *C. elegans* and *Drosophila* oogenesis, the nonhomologous end joining (NHEJ) pathway for DSB repair is actively inhibited early in meiosis (Joyce *et al.* 2012; Lemmens *et al.* 2013; Yin and Smolikove 2013; Lawrence *et al.* 2016; Girard *et al.* 2018) but NHEJ and other pathways, including theta-mediated end joining and single strand annealing, serve as backups to ensure that all DSBs are repaired in late pachytene before the meiotic divisions (Smolikov *et al.* 2007; Macaisne *et al.* 2018). A recent study examining the repair of DNA breaks induced by radiation suggests that mouse spermatocytes switch to a somatic-like repair mode at pachytene, temporarily engaging NHEJ and then HR to repair the damage (Enguita-Marruedo *et al.* 2019). Interestingly, studies in juvenile male mice suggest that structure-specific nucleases may resolve processed DSBs at the expense of the canonical crossover pathway leading to higher levels of meiotic chromosome mis-segregation (Zelazowski *et al.* 2017).

Male meiosis in many species has the added challenge of the presence of heteromorphic sex chromosomes. Meiotic DSBs are induced on hemizygous regions of sex chromosomes (Ashley *et al.* 1995; Moens *et al.* 1997; Sciurano *et al.* 2006; Jaramillo-Lambert and Engebrecht 2010), yet they are unable to participate in crossover formation due to a lack of a homolog. In *C. elegans* and the related nematode, *C. briggsae*, HR using the sister chromatid as repair template, and alternative repair pathways are engaged to repair meiotic DSBs induced on the completely hemizygous X chromosome of males (Checchi *et al.* 2014; Van *et al.* 2016). The presence of hemizygous sex chromosomes also complicates analyses of meiotic recombination in mammals as inactivation of many recombination genes impairs meiotic sex chromosome inactivation (MSCI). MSCI is the process whereby hemizygous regions of sex chromosomes acquire heterochromatin marks and are transcriptionally silenced (Turner 2007). MSCI is required for efficient meiotic progression in males, as failure to inactivate sex chromosomes results in elevated apoptosis and elimination of germ cells (Mahadevaiah *et al.* 2008; Royo *et al.* 2010).

*C. elegans* has an emerged as an excellent model for meiotic studies, including investigations into the sex-specific regulation of meiotic events. Both the *C. elegans* hermaphrodite and male germ lines are arranged in a spatiotemporal gradient that in combination with available molecular markers enables recombination progression to be monitored through all stages of meiotic prophase (Shakes *et al.* 2009; Lui and Colaiacovo 2013; Hillers *et al.* 2015) (Fig 2A). Additionally, the lack of absolute inter-dependence of recombination initiation and chromosome synapsis also facilitates analyses of meiotic mutants. *C. elegans* exists predominantly as a self-fertilizing hermaphrodite (XX); during development, hermaphrodites initially produce sperm and then switch to oocyte production, and thus as adults are functionally female. Males (X0) arise spontaneously due to X chromosome nondisjunction.

**Figure 1.**
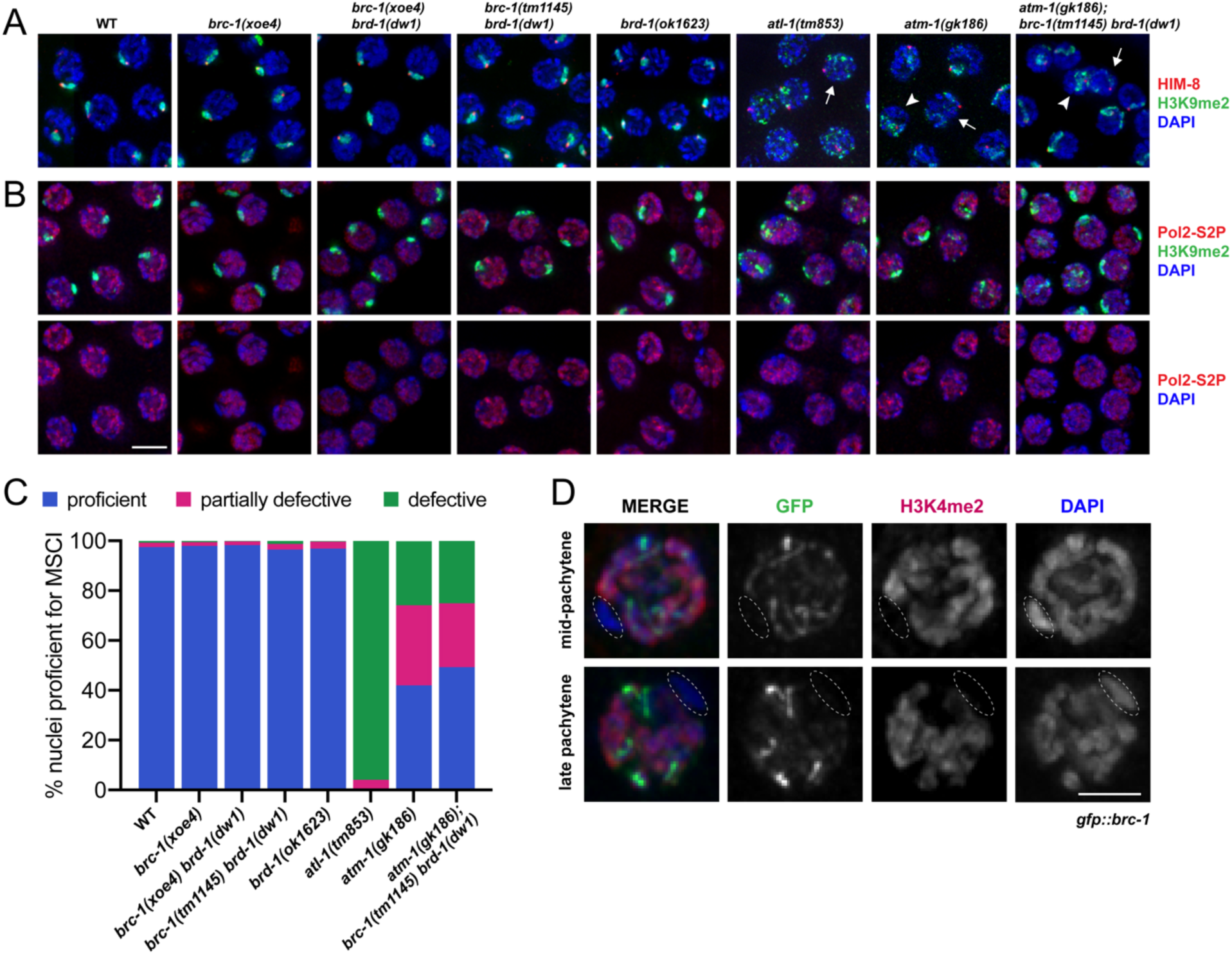
BRC-1-BRD-1 is not required for MSCI. Pachytene nuclei from *C. elegans* wild-type and indicated mutant male germ lines stained with A) anti-H3K9me2 (green; repressive chromatin), anti-HIM-8 (red; X chromosome marker) and counterstained with DAPI (blue) or B) anti-H3K9me2 (green), anti-Pol2-S2P (red; actively transcribing RNA polymerase) and counterstained with DAPI (blue); lower panel shows anti-Pol2-S2P and DAPI. Images are projections through half of the gonad. Scale bar=5μM. C) Quantification of MSCI based on compaction of H3K9me2 and its association with HIM-8; proficient = compact H3K9me2 signal with clear HIM-8 association (blue); partially defective = diffuse H3K9me2 signal associated with HIM-8 (arrowhead in A) (red); defective = H3K9me2 signal spread over the nucleus and no clear HIM-8 association (arrow in A) (green). Number of germ lines, nuclei scored: WT = 3, 433; *brc-1(xoe4)* = 5, 398; *brc-1(xoe4) brd-1(dw1)* = 6, 654; *brc-1(tm1145) brd-1(dw1)* = 3, 257; *brd-1(ok1623)* = 6, 816; *atl-1(tm853)* = 4, 341; *atm-1(gk186)* = 3, 333; *atm-1(gk186); brc-1(tm1145) brd-1(dw1)* = 7, 613. D) GFP::BRC-1 (green) only localizes to synapsed chromosomes and does not localize to the single X chromosome in male meiotic nuclei. X chromosome (circled) identified by chromosome morphology and lack of anti-H3K4me2 staining (red); nuclei counterstained with DAPI (blue). Scale bar=2.5μM.

**Figure 2.**
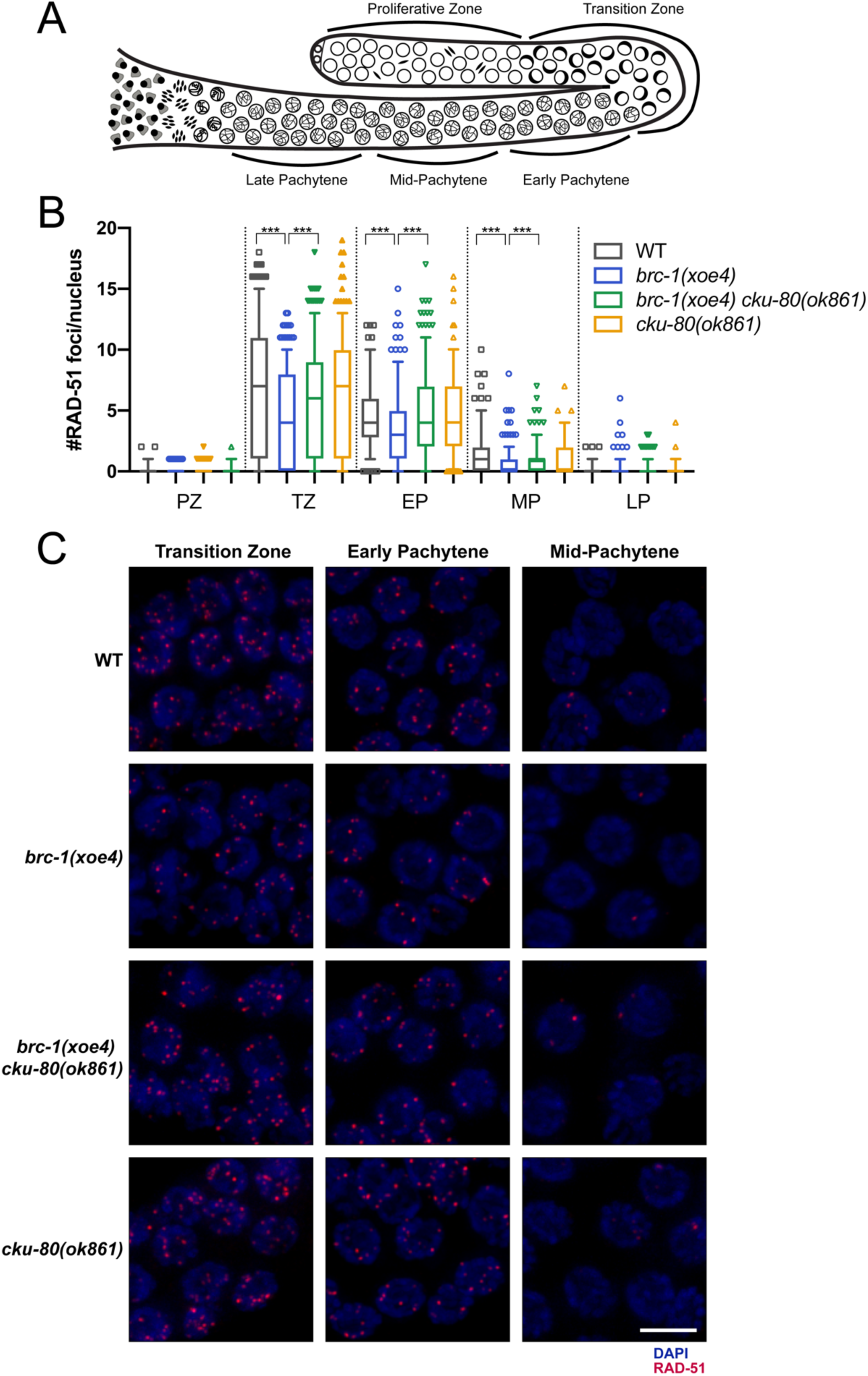
BRC-1 promotes RAD-51 loading in the male germ line. A) Cartoon of the spatiotemporal organization of the *C. elegans* male germ line, modified from Van *et al.* 2016. B) Quantification of RAD-51 in indicated regions of the germ line. Box whisker plots show number of RAD-51 foci per nucleus in the different regions. Horizontal line of each box represents the median, top and bottom of each box represents medians of upper and lower quartiles, lines extending above and below boxes indicate standard deviation and individual data points are outliers from 5–95%. Statistical comparisons by Mann-Whitney of WT versus *brc-1(xoe4)* and *brc-1(xoe4)* versus *brc-1(xoe4) cku-80(ok861)* in the different regions of the germ line; *** p<0.0001. All statistical comparisons are shown in Sup Table 3. PZ =proliferative zone; TZ = transition zone; EP = early pachytene; MP = mid-pachytene; LP = late pachytene. Number of germ lines and nuclei scored in each region: WT = 6, PZ = 958; TZ = 413; EP = 266; MP = 252; LP = 219; *brc-1(xoe4)* = 6, PZ = 848; TZ = 343; EP = 320; MP = 330; LP = 287; *brc-1(xoe4) cku-80(ok861)* = 6, PZ = 905; TZ = 316; EP = 296; MP = 329; LP = 289; *cku-80(ok861)* = 4, PZ = 814; TZ = 287; EP = 202; MP = 230; LP = 217. C) Representative images of nuclei from indicated genotypes and regions of the germ line stained with antibodies against RAD-51 (red) and counterstained with DAPI (blue). Images are projections through half of the gonad. Scale bar=5μm.

The hemizygous X chromosome of *C. elegans* male germ cells undergoes modifications similar to the hemizygous regions of the X and Y of mammalian spermatocytes, including accumulation of repressive chromatin marks resulting in transcriptional silencing (Kelly *et al.* 2002; Reuben and Lin 2002; Bean *et al.* 2004; Maine 2010). A *C. elegans* SETBD1 histone methyltransferase, an ortholog of which has been shown to mediate MSCI in mammals (Hirota *et al.* 2018), and a small RNA pathway are important for silencing the X chromosome of male germ cells (She *et al.* 2009; Bessler *et al.* 2010; Checchi and Engebrecht 2011). However, the role of many components required for MSCI in mammals, including the tumor suppressor E3 ubiquitin ligase BRCA1 and master checkpoint kinase ATR (Turner *et al.* 2004; Royo *et al.* 2013; Broering *et al.* 2014), have not been analyzed in *C. elegans*. Here we examined the requirement for BRCA1-BARD1 (BRC-1-BRD-1) and ATR (ATL-1) in meiotic silencing in *C. elegans*. Surprisingly we found that in contrast to mammals, *C. elegans* BRC-1-BRD-1 is not essential for MSCI. However, MSCI is impaired in the absence of ATL-1, suggesting that while meiotic silencing is conserved, the pathways mediating MSCI have evolved independently. We also find that the meiotic phenotypes of male *brc-1* and *brd-1* mutants are different than those previously reported in female meiosis (Boulton *et al.* 2004; Adamo *et al.* 2008; Janisiw *et al.* 2018; Li *et al.* 2018), providing further evidence that recombination is regulated differently in spermatogenic versus oogenic germ cells (Jaramillo-Lambert and Engebrecht 2010; Checchi *et al.* 2014). Our studies suggest that BRC-1-BRD-1 functions at an early step of meiotic DSB repair in male meiosis, which we propose is similar to one of its established somatic roles in promoting HR at the expense of NHEJ. We also find that this complex alters the crossover landscape when meiosis is perturbed by inhibiting supernumerary crossovers, rather than promoting extra crossovers as in female meiosis. Together, our findings indicate that the processing of meiotic DSBs and the regulation of crossover formation are regulated in a sex-specific manner in *C. elegans*.

## Materials and Methods

### Genetics

*C. elegans* var. Bristol (N2), was used as the wild-type strain. Other strains used in this study are listed in Sup Table 1. Some nematode strains were provided by the Caenorhabditis Genetics Center, which is funded by the National Institutes of Health National Center for Research Resources (NIH NCRR). Strains were maintained at 20°C.

### CRISPR-mediated generation of alleles

*zim-3(xoe15)* was generated in the Bristol background using guides tacgcctgagaacatgtttt and aaaagatcgtgtgatggtcc with repair template: gtaaataacggttgtcgatacgcctgagaacatgtttttggacatttatcttttctagtaggtttttccatatactttattttattctgaa gtttag to delete most of the coding sequence except for exon 7 and 8. External primers cacgacgacaccctcatgta and ttgtgcagagtcgtagcgaa and internal primers cacgacgacaccctcatgta and gctcgtgtacattgagccct were used to genotype for *zim-3(xoe15). brc-1(xoe4)* was introduced into the Hawaiian background (CB4856) using primers, guide and repair template as described (Li *et al.* 2018). *zim-1(xoe6) was* generated in the Bristol and Hawaiian background using guides tccaatcatcacaagtcatc and attcgatgagcttcgtcgtc with repair template tttaaaaatgcagttttaaaagtgtttcattgtcattttatattttccaggcttcgtcgtcgggccgtctgctttttgtaaattgtgtctcatgtgttat to delete the entire coding sequence. External primers cacacatttggctggggtct and atgggcagcagcaagaaagt, and internal primers gctccgtctgcacaaatcct and gttgaaaagcggggaacacc were used to identify *zim-1(xoe6).* Worms were outcrossed a minimum of two times and analyzed phenotypically by examining progeny viability to confirm correct editing.

### Embryonic lethality of male-sired progeny

A single *fog-2(q71)* female was mated with 3 males of indicated genotypes on small *E. coli* OP-50 spots. The mated female was transferred to new plates every 24 hr. Embryonic lethality was determined over 3 days by counting eggs and hatched larvae 24 hr after removing the female and calculating percent as eggs/(eggs + larvae). The progeny of a minimum of 10 mated females were scored.

### Cytological analyses

Immunostaining of germ lines was performed as described (Jaramillo-Lambert *et al.* 2007) except slides were incubated in 100% ethanol instead of 100% methanol for direct GFP fluorescence of GFP::COSA-1. The following primary antibodies were used at the indicated dilutions: rabbit anti-Pol2 Ser2-P (1:500; cat #ab5059; Abcam, Cambridge, MA; RRID: AB_304749), rabbit anti-HIM-8 (1:500; cat #4198.00.02; SDIX; Newark, DE; RRID: AB_2616418), rabbit anti-histone H3K4me2 (1:500; cat# 9725; Cell Signaling Technology; Danvers, MA; RRID: AB_10205451), mouse anti-histone H3K9me2 1:500 (1:500; Cat# 9753; AbCam; Cambridge, MA; RRID: AB_659848), rabbit anti-RAD-51 (1:10,000; cat #2948.00.02; SDIX; RRID: AB_2616441), mouse anti-GFP (1:500; cat #632373; BD Biosciences; San Jose, CA). Secondary antibodies Alexa Fluor 594 donkey anti-rabbit IgG, Alexa Fluor 594 goat anti-mouse IgG, Alexa Fluor 488 goat anti-rabbit IgG and Alexa Fluor 488 goat anti-mouse IgG from Life Technologies were used at 1:500 dilutions. DAPI (2μg/ml; Sigma-Aldrich) was used to counterstain DNA.

Collection of fixed images was performed using an API Delta Vision deconvolution microscope or an API Delta Vision Ultra equipped with an 60x, NA 1.49 objective lens, and appropriate filters for epi-fluorescence. Z stacks (0.2 μm) were collected from the entire gonad. A minimum of three germ lines was examined for each condition. Images were deconvolved using Applied Precision SoftWoRx batch deconvolution software and subsequently processed and analyzed using Fiji (ImageJ) (Wayne Rasband, NIH).

RAD-51 foci were quantified in a minimum of three germ lines of age-matched males (18-24 hr post-L4). We divided germ lines into the transition zone (leptotene/zygotene), as counted from the first and last row with two or more crescent-shaped nuclei, and then divided pachytene into 3 equal parts: early, mid and late (Fig 2A). RAD-51 were quantified from half projections of the germ lines. The number of foci per nucleus was scored for each region.

To assess formation of RAD-51 foci following IR treatment, 18-24 hr post-L4 male worms were exposed to 10 Grays (Gys) of ionizing radiation (IR); 1 hr post IR, worms were dissected and gonads fixed for immunofluorescence as above.

GFP::COSA-1 foci were quantified from deconvolved 3D data stacks; late pachytene nuclei were scored individually through z-stacks to ensure that all foci within each individual nucleus were counted.

For live cell imaging, 18-24 hr post L4 males were anesthetized in 1mM tetramisole (Sigma-Aldrich) and immobilized between a coverslip and an 2.5% agarose pad on a glass slide. Z-stacks (0.33 μm) were captured on a spinning-disk module of an inverted objective fluorescence microscope [Marianas spinning-disk confocal (SDC) real-time 3D Confocal-TIRF (total internal reflection) microscope; Intelligent Imaging Innovations] with a 100x, 1.46 numerical aperture objective, and a Photometrics QuantiEM electron multiplying charge-coupled device (EMCCD) camera. Z-projections of approximately 20-30 z-slices were generated, cropped, and adjusted for brightness in ImageJ.

### Meiotic mapping

Meiotic crossover frequencies and distribution were assayed using single-nucleotide polymorphism (SNP) markers as in (Nabeshima *et al.* 2004). The SNP markers located at the boundaries of the chromosome domains were chosen based on data from WormBase (WS231) and (Bazan and Hillers 2011; Saito *et al.* 2013). Markers and primers used are listed in Sup Table 2. Hawaiian strain CB4856 males carrying each mutation were crossed to the same mutant strain in the Bristol background. Among the progeny of this cross, male worms were plated individually and crossed to two *fog-2(q71)* female worms in the Bristol background. Upon successful mating, embryos (Smolikov *et al.* 2008) together with larva up to L4 stage were collected individually and stored at - 80°C. Since all three mutant (*brc-1, zim-1, brc-1;zim-1*) hermaphrodites produce self-fertilized male progeny, the identity of the hybrid Bristol/Hawaiian male was confirmed by PCR and restriction digest before the collected samples were used for further analysis: individuals were lysed in 5μl of lysis buffer (50 mM KCl, 10 mM Tris pH8.2, 2.5 mM MgCl2, 0.45% NP-40, 0.45% Tween20, 0.01% gelatin; 60μg of proteinase K/ml was added before use) and diluted to 50μl volume with molecular biology grade water. PCR was performed using 3-5μl diluted lysate with Phusion or Taq polymerase in a 15μl reaction. Half volume of the PCR products was digested overnight with appropriate restriction enzyme and analyzed on 1%-2.5% agarose gels. Confirmation of double crossovers were performed either with additional SNPs by a distinctive restriction enzyme digest or by repeating PCR and digestion if no additional SNPs were available for the marker as described in (Saito *et al.* 2013) (Sup Table 2).

### Statistical analyses

Statistical analyses and figures were prepared using GraphPad Prism version 8.0 (GraphPad Software). RAD-51 foci numbers were analyzed by Mann-Whitney. Fisher exact test on a 2-by-2 contingency table was used for statistical analyses on genetic map distance and distribution. For statistical analyses of interference, χ2 tests on 2-by-2 contingency tables of observed and expected DCOs were performed (Brady *et al.* 2018). Detailed descriptions of statistical analyses are indicated in figure legends.

### Data availability

Strains and reagents are available upon request. The authors affirm that all data necessary for confirming the conclusions of this article are represented fully within the article and its tables and figures. Supplemental data are deposited at figshare.

## Results

### *C. elegans* BRC-1-BRD-1 is not required for MSCI

During *C. elegans* meiosis, the X chromosome accumulates the repressive chromatin mark histone H3 lysine 9 di-methylation (H3K9me2) and is transcriptionally silenced similar to MSCI in mammals (Kelly *et al.* 2002; Reuben and Lin 2002; Bean *et al.* 2004; Checchi and Engebrecht 2011). In mice, the E3 ubiquitin ligase BRCA1, critical for DNA damage response, is essential for MSCI. As a result, *brca1*^*-/-*^ mutant male germ cells inappropriately express X-linked genes leading to pachytene arrest, apoptosis of spermatocytes and infertility (Xu *et al.* 2003; Turner *et al.* 2004; Broering *et al.* 2014). To determine whether *C. elegans* BRC-1 or its binding partner BRD-1 (Boulton *et al.* 2004) plays a role in MSCI, we stained male *brc-1, brd-1* and *brc-1 brd-1* double mutant germ lines [*brc-1(xoe4), brd-1(ok1623), brc-1(xoe4) brd-1(dw1)* and *brc-1(tm1145) brd-1(dw1)* (Polanowska *et al.* 2006; Janisiw *et al.* 2018; LI *et al.* 2018)] with antibodies against H3K9me2 and the X-specific pairing center binding protein HIM-8 (Phillips and Dernburg 2006). The X chromosome, marked by HIM-8, was highly enriched for H3K9me2 in all of the *brc-1* and *brd-1* mutant combinations, as in wild type, suggesting that enrichment of this repressive chromatin mark on the X chromosome occurs in the absence of BRC-1 and/or BRD-1 (Fig. 1A, C). To examine the transcriptional status of the X chromosome, we co-stained germ lines with antibodies that recognize H3K9me2 and RNA polymerase II phosphorylated on serine 2 (Pol2-S2P), which is associated with transcriptional elongation (Hsin and Manley 2012), and for which we previously showed is excluded from the single X chromosome in male germ cells (Larson *et al.* 2016). Pol2-S2P was present throughout the nucleus except for a single chromosome, marked by H3K9me2, in all *brc-1/brd-1* mutants (Fig. 1B), suggesting that the X chromosome is transcriptionally silenced in the absence of BRC-1-BRD-1.

In mammals, BRCA1 is observed on asynapsed axes and is enriched on the X-Y sex body (Turner *et al.* 2004). In *C. elegans* hermaphrodites, BRC-1 and BRD-1 become associated with fully synapsed chromosomes in pachytene (Polanowska *et al.* 2006; Janisiw *et al.* 2018; Li *et al.* 2018). We examined the localization of BRC-1 in male germ lines expressing an endogenously tagged and fully functional GFP fusion [GFP::BRC-1; (Li *et al.* 2018)] and found that it was also associated with tracks corresponding to synapsed chromosomes at pachytene. However, in contrast to the six tracks observed in oocytes, only five tracks were present in spermatocytes, suggesting that BRC-1-BRD-1 does not localize to the asynapsed X chromosome. To verify this, we co-stained male germ lines with antibodies against GFP, to detect GFP::BRC-1, and the activating chromatin mark, H3K4me2, which is enriched on all chromosomes except the X (Reuben AND LIN 2002; Bean *et al.* 2004; Jaramillo-Lambert and Engebrecht 2010; Checchi and Engebrecht 2011), and found that the chromosome lacking H3K4me2 also lacked GFP::BRC-1 (Fig. 1D). Thus, contrary to mammals, *C. elegans* BRC-1-BRD-1 is not enriched on asynapsed sex chromosomes in male germ cells.

During mammalian MSCI, BRCA1 facilitates the recruitment of the Phosphoinositide 3-kinase ataxia telangiectasia and RAD3-related (ATR) kinase to sex chromosomes; ATR in turn phosphorylates the histone variant H2AX (γ-H2AX) to facilitate chromosome compaction. Consequently, inactivation of either ATR or H2AX also results in MSCI failure (Fernandez-Capetillo *et al.* 2003; Turner *et al.* 2004; Royo *et al.* 2013). Given that BRC-1-BRD-1 is not essential for MSCI and no H2AX variant has been identified in the *C. elegans* genome (Boulton 2006), we next addressed whether ATL-1 is required for MSCI. To that end, we monitored the localization of H3K9me2 and HIM-8 in *atl-1(tm853)* deletion mutant germ lines. In contrast to *brc-1* or *brd-1* mutants, mutation of *atl-1* resulted in altered distribution of H3K9me2. In most nuclei (95.9%), there was no clear association between HIM-8 and H3K9me2, indicating that the X chromosome was not specifically enriched for H3K9me2, and in the remaining nuclei (4.1%), H3K9me2 was associated with HIM-8 but had a much less compact signal (Fig. 1A, C). Analysis of Pol2-S2P also showed that the X chromosome was not transcriptionally silenced, suggesting that the absence of ATL-1 results in MSCI failure (Fig. 1B). Thus, although BRC-1-BRD-1 does not appear to play a role in MSCI, ATL-1 is essential for the correct targeting of H3K9me2 and silencing of the X chromosome during *C. elegans* male meiosis.

ATR participates with the related and partially redundant kinase, ataxia-telangiectasia mutated (ATM) during DNA damage signaling (Abraham 2001). In mice, ATM does not play a role in MSCI (Royo *et al.* 2013). To determine whether ATM functions in MSCI in *elegans*, we monitored H3K9me2 and HIM-8 in germ lines of the *atm-1(gk186)* deletion mutant. While 42.1% of nuclei were wild type with respect to clear and compact association between HIM-8 and H3K9me2, 32.1% of nuclei showed association between the signals but much more diffuse H3K9me2 staining, and 25.8% showed no association between HIM-8 and H3K9me2 (Fig. 1A, C). Similarly, Pol2-S2P showed a variable staining pattern with some nuclei containing a single chromosome lacking Pol2-S2P and enriched for H3K9me2, which presumably corresponds to the X chromosome, while in other nuclei no clear chromosome lacking Pol2-S2P was detected (Fig. 1B). Thus, in *C. elegans*, ATL-1, and to a lesser extent ATM-1, are important for MSCI.

To determine whether a function of BRC-1-BRD-1 in MSCI can be uncovered in the sensitized *atm-1* mutant background, we examined H3K9me2 and Pol2-S2P in the *atm-1(gk186); brc-1(tm1145) brd-1(dw1)* triple mutant. We found no difference in either H3K9me2 or Pol2-S2P localization between *atm-1(gk186)* and *atm-1(gk186); brc-1(tm1145) brd-1(dw1)*, consistent with BRC-1-BRD-1 being dispensable for MSCI in *C. elegans*.

### A subset of meiotic DSBs is repaired by NHEJ in the absence of BRC-1-BRD-1 in male germ cells

BRCA1-BARD1 has been implicated in promoting HR repair in somatic cells; however, its role in meiotic recombination has been controversial and is complicated by the pachytene arrest and apoptotic removal of *brca1* mutant spermatocytes due to MSCI failure (Xu *et al.* 2003; Broering *et al.* 2014). The finding that neither *brc-1* nor *brd-1* mutants impair MSCI in *C. elegans* prompted us to examine the role of BRC-1-BRD-1 in meiotic recombination in the absence of the complications associated with MSCI failure. To that end, we monitored meiotic DSB repair by examining the assembly and disassembly of the recombinase RAD-51 (Rinaldo *et al.* 2002) in the spatiotemporal organization of the *C. elegans* male germ line (Colaiacovo *et al.* 2003; Checchi *et al.* 2014) (Fig. 2A).

*brc-1* and *brd-1* mutant hermaphrodites exhibit a slight increase in embryonic lethality and male progeny (a readout of X chromosome nondisjunction), and some RAD-51 foci perdure in late meiotic prophase, suggesting that repair of a subset of meiotic DSBs is delayed in the absence of BRC-1-BRD-1 (Boulton *et al.* 2004; Adamo *et al.* 2008; Janisiw *et al.* 2018; Li *et al.* 2018). In contrast to the appearance of more RAD-51 foci in mid and late pachytene in female germ cells, fewer RAD-51 foci were observed in *brc-1, brd-1* or *brc-1 brd-1* male germ cells compared to wild type in early meiotic prophase (transition zone) through mid-pachytene (Fig. 2B, C; Sup Fig 1). These results suggest that in the absence of BRC-1-BRD-1 either fewer DSBs are induced, a subset of DSBs is repaired without loading RAD-51, or repair occurs with faster kinetics than wild type. Given a role of BRCA1 in promoting HR at the expense of NHEJ in somatic cells (Daley and Sung 2014), we tested the hypothesis that some meiotic DSBs are repaired by NHEJ in the absence of BRC-1-BRD-1 in male germ cells. To that end, we simultaneously inactivated BRC-1 or BRD-1 and CKU-80 or CKU-70, the *C. elegans* KU80/KU70 orthologs that mediate NHEJ and monitored RAD-51 foci throughout the germ line (Fig 2B, C; Sup Fig 1). When NHEJ was inactivated in the *brc-1* or *brd-1* mutants, RAD-51 foci were restored to wild-type levels in transition zone through mid-pachytene in male germ cells. We also observed a small, but statistically significant elevation of RAD-51 in late pachytene when both BRC-1-BRD-1 and NHEJ were removed, suggesting that both of these complexes contribute to repair of lesions at late pachytene (Sup Fig 1B; Sup Table 3), similar to what has been observed in oogenesis (Smolikov *et al.* 2007; Adamo *et al.* 2008). Together, these results suggest that BRC-1-BRD-1 functions at an early step of DSB processing to facilitate repair by HR in male germ cells similar to its proposed role in somatic cells, and in its absence, some breaks are channeled through NHEJ in early meiotic prophase.

### GFP::BRC-1 concentration at foci in early meiotic prophase is dependent on resected DSBs

We next examined the localization of GFP::BRC-1 by live cell imaging. In wild-type male germ lines, GFP::BRC-1 was nucleoplasmic and formed a small number of bright foci in proliferating germ cells (Fig 3A, B). As cells progressed into meiosis, GFP::BRC-1 was observed in multiple foci at transition zone and early pachytene; tracks of fluorescence were also beginning to form at early pachytene (Fig 3A, B). At mid-pachytene, GFP::BRC-1 was predominantly in tracks, which had begun to concentrate on a chromosomal subdomain. Further concentration into 5 stretches and then puncta, were observed in late pachytene through diplotene. The dynamic localization of GFP::BRC-1 in the male germ line is similar to the hermaphroditic germ line: GFP::BRC-1 foci partially overlap with RAD-51 (Sup Fig 2A), suggesting they mark sites of ongoing meiotic recombination, and the GFP::BRC-1 tracks in pachytene colocalize with the synaptonemal complex (SC) that become concentrated on the short arm dependent on crossover formation (Li *et al.* 2018).

**Figure 3.**
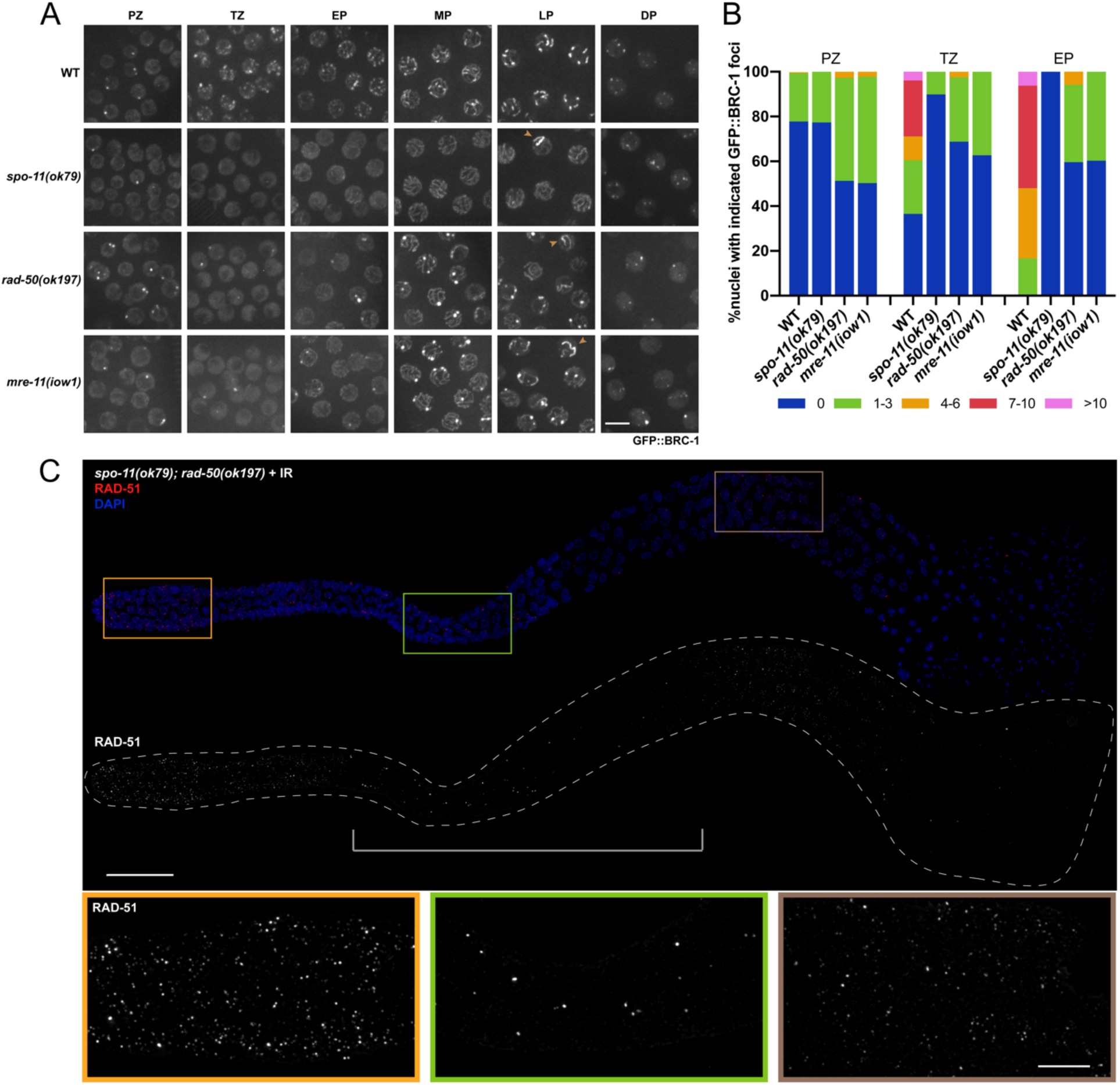
GFP::BRC-1 concentration at foci in early meiotic prophase is dependent on meiotic DSB resection. A) High-magnification images of live worms expressing GFP::BRC-1 from the indicated genetic backgrounds and gonad region (PZ = proliferative zone, TZ = transition zone, EP = early pachytene, MP = mid-pachytene, LP = late pachytene, DP = diplotene). Images are projections through half of the gonad. Scale bar=5μm. B) Number of GFP::BRC-1 foci in PZ, TZ, and EP in wild type and mutants. Numbers were binned as 0, 1-3, 4-6, 7-10, >10. A minimum of 3 germ lines were quantified for each genotype. C) *spo-11(ok79); rad-50(ok197)* male gonad fixed and dissected 1 h after exposure to 10 Greys IR, stained with anti-RAD- 51 antibody (red) and counterstained with DAPI (blue). In the *spo-11; rad-50* mutant RAD-51 foci are largely absent in most nuclei in the central portion of the gonad, indicated by the bracket, from the onset of meiotic prophase to mid-pachytene. Images are projections through the entire gonad. Four germ lines were examined. Scale bar=20μm. Insets show selected nuclei from different regions of the germ line; scale bar=5μm.

To test the dependencies of BRC-1 localization on DSB formation and processing, we examined GFP::BRC-1 in *spo-11, rad-50 and mre-11* mutants. *spo-11* mutants are unable to form meiotic DSBs (Dernburg *et al.* 1998), while *rad-50* and *mre-11* are required for both DSB formation and processing (Chin and Villeneuve 2001; Hayashi *et al.* 2007). In the absence of SPO-11, very few GFP::BRC-1 foci were present in transition zone and early pachytene compared to wild type (Fig 3A, B). At early to mid-pachytene GFP::BRC-1 was observed in tracks in the *spo-11* mutant similar to wild type, as synapsis occurs in the absence of genetic recombination in *C. elegans* (Dernburg *et al.* 1998) (Fig 3A). In late pachytene, GFP::BRC-1 fluorescence did not concentrate on a portion of each chromosome pair as in wild type, consistent with these events being dependent on crossover formation. However, in 10.7±3.2% of pachytene nuclei there was enrichment of GFP::BRC-1 on an entire chromosome (Fig 3A, arrowhead) with weak fluorescence on the other synapsed chromosomes. This has been observed for GFP::BRC-1 and other synapsis markers in oogenesis and likely represents *spo-11*-independent lesions capable of recruiting meiotic DNA repair components and altering SC properties (Machovina *et al.* 2016; Nadarajan *et al.* 2017; Pattabiraman *et al.* 2017; Li *et al.* 2018).

We next examined the requirement for RAD-50 and MRE-11 in recruitment of GFP::BRC-1 to early meiotic foci. RAD-50 and MRE-11 form a complex with NBS-1 (MRX/N complex) and are required for both DSB formation and processing for repair through HR in meiotic cells, in addition to playing a role in repair of lesions generated during DNA replication (Chin and Villeneuve 2001; Hayashi *et al.* 2007; Girard *et al.* 2018). In *rad-50(ok197)* and *mre-11(ok179)* null mutants, GFP::BRC-1 was observed in fewer foci compared to wild type in transition zone and early pachytene (Fig 3A, B, Sup Fig 2B). However, in contrast to *spo-11*, an increased number of nuclei with 1-3 GFP::BRC-1 foci were present in proliferating germ cells and throughout meiotic prophase (Fig. 3A, B), suggesting GFP::BRC-1 is enriched at lesions generated during S phase in the these mutant backgrounds. We also observed an earlier appearance and higher percentage of nuclei showing concentrated signal on a subset of chromosomes (*rad-50(ok197)*, 21.17±4.6%), consistent with recruitment of recombination proteins and alteration of the SC properties at mitotic lesions as they progress through meiosis. Together, these results suggest that the enrichment of GFP::BRC-1 to abundant foci in early meiotic prophase is dependent on meiotic DSB formation.

To determine the requirement for DSB end processing in recruiting GFP::BRC-1 to sites of meiotic recombination, we took advantage of a separation-of-function allele, *mre-11(iow1);* worms harboring this allele are competent for meiotic DSB formation but defective in resection (Yin and Smolikove 2013). As with *rad-50(ok197)* and *mre-11(ok179)* null mutants, there was a reduction in meiotic GFP::BRC-1 foci in *mre-11(iow1)* mutant germ lines (Fig 3A, B) and a similar number of pachytene nuclei showing concentration of GFP::BRC-1 on a subset of chromosomes (*mre-11(iow1)*, 19.23±2.8%). These results suggest that concentration of GFP::BRC-1 into foci in early meiotic prophase requires DSB resection, consistent with BRC-1-BRD-1 functioning at an early step of meiotic DSB processing to promote HR at the expense of NHEJ.

### RAD-51 loading is dependent on RAD-50 in male meiotic germ cells

Meiotic recombination occurs in the context of specialized chromosome structure, the chromosomal axes and fully formed SC, to promote interhomolog crossovers. Previous analyses in oogenic germ lines revealed a requirement for RAD-50 in loading RAD-51 at DSBs in meiotic prophase (Hayashi *et al.* 2007). Given the somatic-like role of BRC-1-BRD-1 in promoting HR at the expense of NHEJ in meiotic male germ cells, we next addressed whether male meiosis also requires RAD-50 for loading RAD-51 in the context of synapsed chromosomes. To that end, we analyzed RAD-51 localization in *rad-50; spo-11* males. The *spo-11* mutation was introduced to eliminate all endogenous meiotic DSBs. DNA breaks were induced by 10 Gys of IR and 1 hr post IR, gonads were dissected and stained with antibodies against RAD-51 (Hayashi *et al.* 2007). Similar to hermaphroditic germ lines, abundant IR-induced RAD-51 foci were observed in proliferating germ cells and in mid-late pachytene/diplotene spermatocytes (Fig 3C). However, irradiated *spo-11; rad-50* double mutant germ lines contained a region extending from the transition zone to mid-late pachytene in which very few foci were observed. Thus, similar to oogenesis, RAD-51 loading is dependent on RAD-50 during meiotic prophase in spermatogenic germ lines. Together, our genetic and cell biological analyses of BRC-1-BRD-1 and DSB processing factors suggest that properties of both somatic and meiotic repair modes exist in male germ cells.

### BRC-1-BRD-1 is important when crossover formation is blocked on a subset of chromosomes during spermatogenesis

In somatic cells, BRCA1 plays a critical role when errors in the cell cycle occur (Takaoka and Miki 2018) and we previously found that removal of BRC-1-BRD-1 during oogenesis impairs progeny viability and RAD-51 stabilization when crossover formation is blocked on a subset of chromosomes (Li *et al.* 2018). To examine the consequence of inactivating BRC-1-BRD-1 under similar conditions during male meiosis, we monitored the viability of progeny sired by mutant *zim-1(tm1813)* [chromosomes *II* and *III* fail to pair and synapse (Phillips and Dernburg 2006)], *brc-1(xoe4); zim-1(tm1813), brc-1(tm1145); zim-1(tm1813)* and *brd-1(ok1623); zim-1(tm1318)* males. *brc-1(tm1145)* is a hypo-morphic allele that we previously showed impairs recombination under meiotic checkpoint activating conditions in oogenesis (Li *et al.* 2018). We used worms carrying the *fog-2(q71)* mutation for these experiments to eliminate hermaphrodite spermatogenesis, rendering XX animals self-sterile (Schedl and Kimble 1988), so that the contribution of the male parent to embryonic lethality could be assessed unambiguously. Similar to our findings in hermaphrodites (Li *et al.* 2018), removal of BRC-1 or BRD-1 enhanced the embryonic lethality of *zim-1* mutants when mutant sperm were used to fertilize *fog-2* ova (Fig.4A; p<0.0001 by One Way ANOVA). These results suggest that BRC-1-BRD-1 plays important roles to enhance the quality of male germ cells under meiotic checkpoint activating conditions.

Previous analyses in the hermaphrodite germ line revealed that RAD-51 levels are elevated genome wide when the obligate crossover is not established on any or all chromosome pairs (Colaiacovo *et al.* 2003; Carlton *et al.* 2006; Mets and Meyer 2009). Removal of BRC-1-BRD-1 under these conditions resulted in a “dark zone” of RAD-51 in mid-late pachytene, which is likely a consequence of premature RAD-51 disassembly (Li *et al.* 2018). To determine whether BRC-1-BRD-1 promotes RAD-51 filament stability in male germ lines when not all chromosomes are connected by a crossover, we monitored RAD-51 levels in *zim-1* mutants in the presence and absence of BRC-1-BRD-1. Similar to oogenic germ lines, blocking crossover formation on a subset of chromosomes resulted in elevated levels of RAD-51 foci throughout meiotic prophase in male germ lines (Fig 4 B, C). However, in the absence of BRC-1, we did not observe a RAD-51 “dark zone”, suggesting that BRC-1-BRD-1 does not play a role in stabilizing the RAD-51 filament under checkpoint activating conditions in male germ cells (Fig 4C). Quantification of foci revealed reduced RAD-51 levels in *brc-1; zim-1* compared to *zim-1* (Fig. 4B), similar to the reduction in RAD-51 foci observed in *brc-1* or *brd-1* mutants alone compared to wild-type males (Fig. 2B). However, the RAD-51 levels in *brc-1; zim-1* are still significantly higher throughout pachytene than in wild-type male germ lines (compare Fig 2A and Fig 4B). These results suggest that BRC-1-BRD-1 promotes meiotic recombination in spermatogenesis using different mechanisms than in oogenesis under meiotic checkpoint activation.

**Figure 4.**
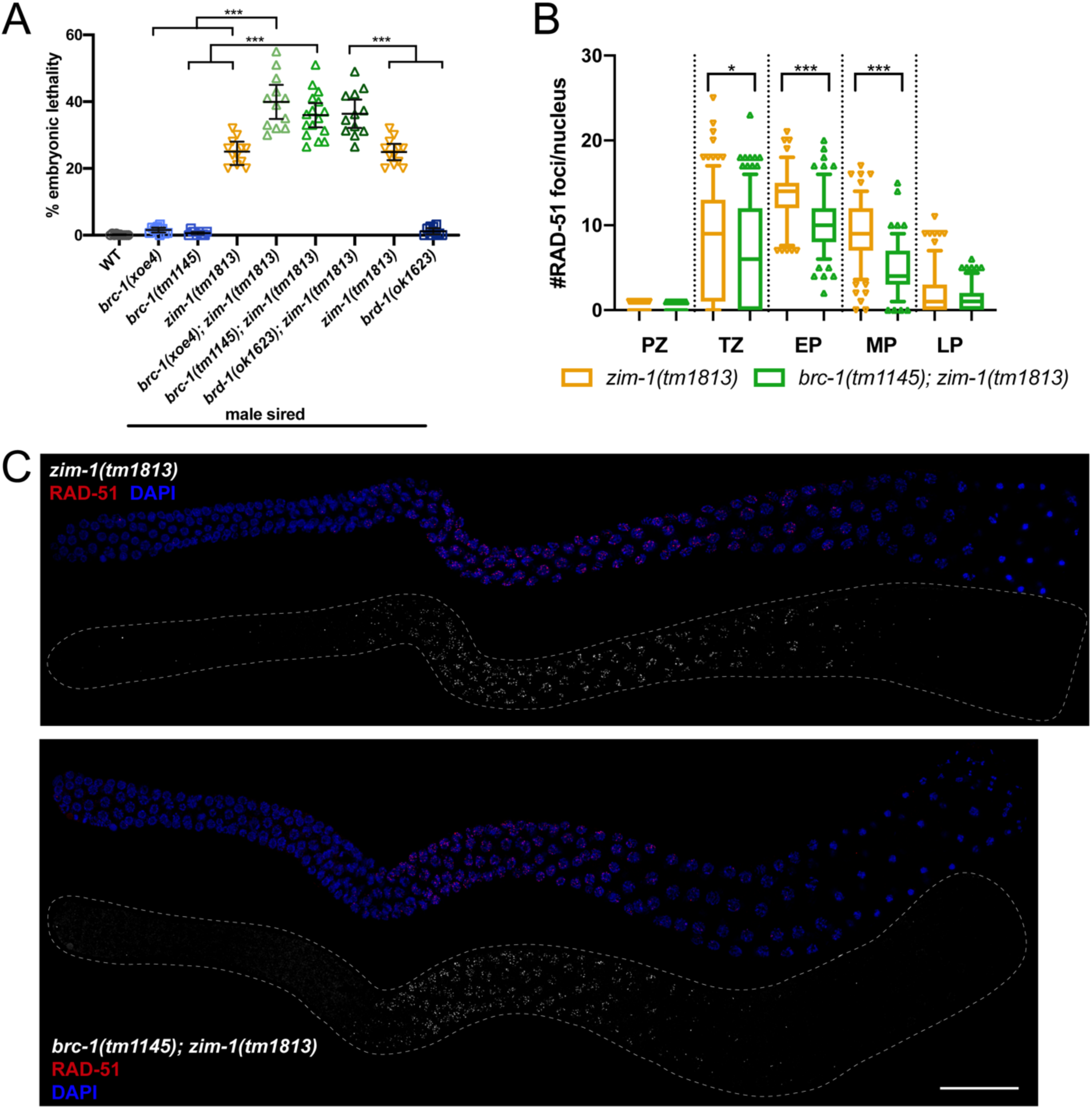
Progeny embryonic lethality is enhanced when sired by *brc-1/brd-1*; *zim-1* double mutant males but RAD-51 stability in not impaired. A) Embryonic lethality of *fog-2(q71)* progeny sired by *brc-1(xoe4), brc-1(tm1145), zim-1(tm1813), brc-1(xoe4); zim-1(tm1813), brc-1(tm1145); zim-1(tm1813), brd-1(ok1623); zim-1(tm1813), brd-1(ok1623)* males. Mean and 95% confidence intervals are shown. The genetic interaction between *brc-1/brd-1* and *zim-1* is significant by a one-way ANOVA (***p<0.0001). A minimum of 10 worms were scored for each genotype. B) Box whisker plots show average number of RAD-51 foci per nucleus in the different zones. Horizontal line of each box indicates the median, the top and bottom of the box indicates medians of upper and lower quartiles, lines extending above and below boxes indicate standard deviation and individual data points are outliers from 5-95%. Statistical comparisons by Mann-Whitney of *zim-1(tm1813)* versus *brc-1(tm1145); zim-1(tm1813)* in the different regions of the germ line: * p<0.05; *** p<0.0001. PZ =proliferative zone; TZ = transition zone; EP = early pachytene; MP = mid-pachytene; LP = late pachytene. Numbers of nuclei scored from 4 germ lines in each zone for *zim-1*: PZ = 668; TZ = 237; EP = 111; MP = 151; LP – 167 and *brc-1; zim-1*: PZ = 545; TZ = 318; EP = 155; MP = 137; LP = 149. C) *zim-1(tm1813)* and *brc-1(tm1145); zim-1(tm1813)* mutant germ lines stained with anti-RAD-51 antibody (red) and counterstained with DAPI (blue). Images are projections through half of the gonad. A minimum of 4 germ lines were imaged. Scale bar=20μm.

### BRC-1-BRD-1 inhibits COSA-1 marked crossover designation when meiosis is perturbed in male germ cells

In addition to stabilizing the RAD-51 filament, BRC-1-BRD-1 promotes formation of crossover precursors marked by the cyclin related COSA-1 (Yokoo *et al.* 2012) in the *zim-1* mutant background in hermaphrodites (Li *et al.* 2018). To determine whether BRC-1-BRD-1 influences crossover designation in male germ cells, we monitored GFP::COSA-1 (Yokoo *et al.* 2012) in *brc-1, brd-1, zim-1, brc-1; zim-1* and *brd-1; zim-1* mutant germ lines. Wild-type males mostly exhibit five COSA-1 foci, one on each of the five pairs of autosomes but not on the single X chromosome (Checchi *et al.* 2014). This pattern was unaltered by removal of either BRC-1 or BRD-1 (WT = 4.99± 0.30; *brc-1(xoe4)* = 4.99±0.30; *brd-1(ok1623)* = 5.02±0.28; Fig. 5A). As *zim-1* mutants have two asynapsed chromosome pairs, we expected to observe three COSA-1 foci; however, we observed an average of 4.61±1.12 COSA-1 foci (Fig. 5A). Further, removing BRC-1 or BRD-1 in *zim-1* males resulted in more COSA-1 foci (*brc-1(xoe4); zim-1(tm1813)* = 5.32±0.97; *brd-1(ok1623); zim-1(tm1813)* = 5.29±0.99) (Fig. 5A, B). This is opposite to what we observed in hermaphrodites, where reduced levels of GFP::COSA-1 was observed in the absence of BRC-1 or BRD-1 in *zim-1* mutants (Li *et al.* 2018).

**Figure 5.**
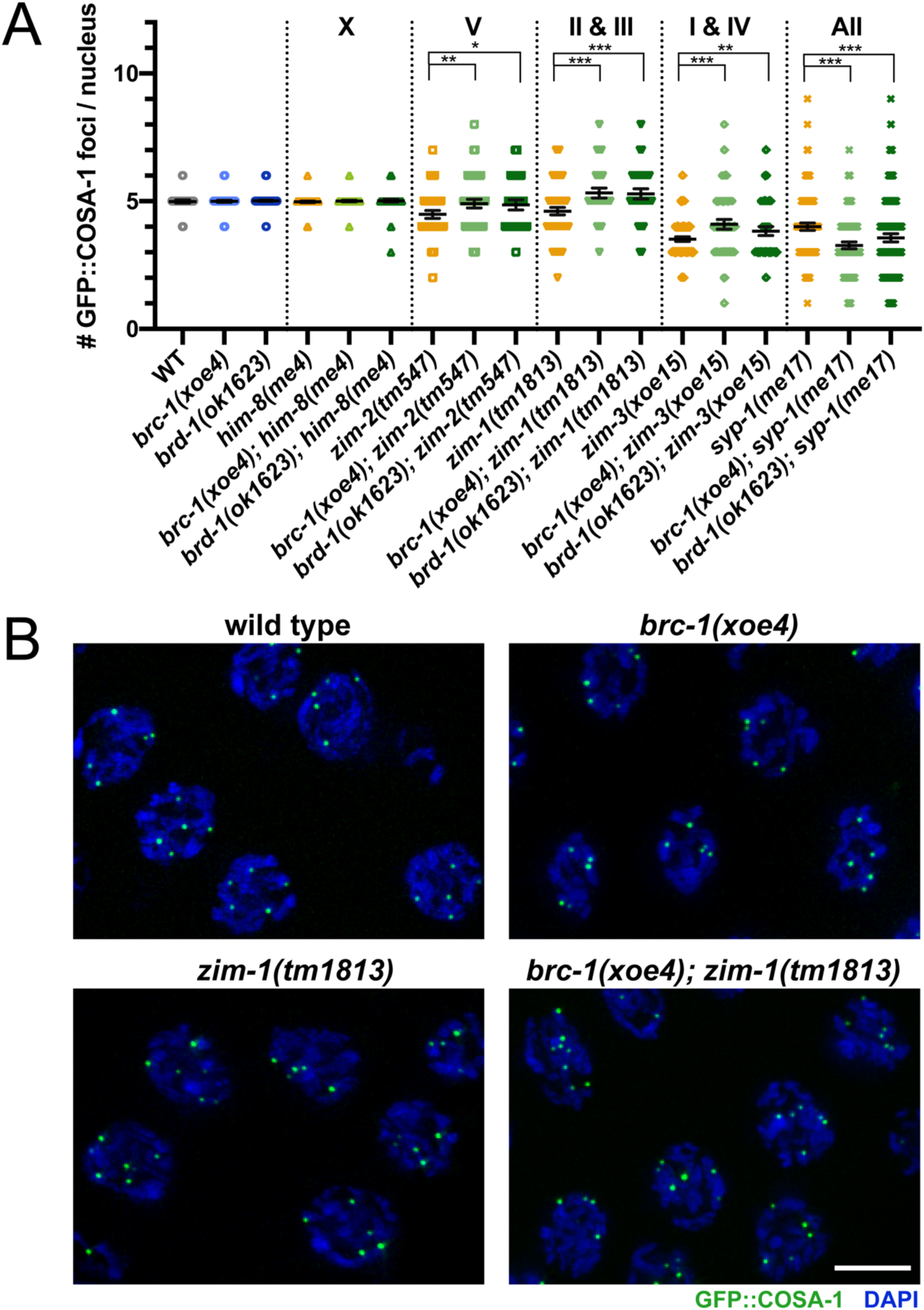
BRC-1-BRD-1 inhibits GFP::COSA-1 marked crossover precursors when a subset of chromosomes fail to form crossovers. A) Number of COSA-1 foci in mid-late pachytene in indicated mutants; mean and 95% confidence intervals are shown. Number of nuclei scored: *gfp::cosa-1* = 97, *gfp::cosa-1; brc-1(xoe4)* = 194, *gfp::cosa-1; brd-1(ok1623)* = 103, *gfp::cosa-1; him-8(me4)* = 151, *gfp::cosa-1; brc-1(xoe4); him-8(me4)* = 183; *gfp::cosa-1; brd-1(ok1623); him-8(me4)* = 172, *gfp::cosa-1; zim-2(tm547)* = 125, *gfp::cosa-1; brc-1(xoe4); zim-2(tm547)* = 128; *gfp::cosa-1; brd-1(ok1623); zim-2* = 84, *gfp::cosa-1; zim-1(tm1813)* = 120, *gfp::cosa-1*; *brc-1(xoe4); zim-1(tm1813)* = 100, *gfp::cosa-1; brd-1(ok1623); zim-1(tm1813)* = 97, *gfp::cosa-1; zim-3(xoe15)* = 308, *gfp::cosa-1*; *brc-1(xoe4); zim-3(xoe15)* = 133, *gfp::cosa-1; brd-1(ok1623); zim-3(xoe15)* = 145, *gfp::cosa-1; syp-1(me17)* = 271, *gfp::cosa-1*; *brc-1(xoe4); syp-1(me17)* = 281, *gfp::cosa-1; brd-1(ok1623); syp-1(me17)* = 344. B) Half projections of late pachytene region showing GFP::COSA-1 (green) and DAPI (blue) in wild type, *brc-1(xoe4), zim-1(tm1813)* and *brc-1(xoe4); zim-1(tm1813)*. Scale bar=5μm.

To examine this further, we monitored GFP::COSA-1 foci in additional mutants that lead to asynapsis of different chromosome pairs. Pairing and synapsis of the *X* chromosome is impaired in *him-8* mutants, *zim-2* mutants have asynapsed chromosome *Vs* and two chromosome pairs, *I* and *IV*, fail to pair and synapse in *zim-3* mutants (Phillips *et al.* 2005; Phillips AND DERNBURG 2006). As expected, mutation of *him-8* had no effect on GFP::COSA-1 levels either in the presence or absence of BRC-1-BRD-1, presumably due to the presence of the single X chromosome in male germ cells (Fig. 5A). *zim-2* and *zim-3* mutants showed higher than expected average of COSA-1 foci (*zim-2(tm547)*=4.48±0.85 observed vs 4 expected, *zim-3(xoe15)*=3.52±0.80 observed vs 3 expected), similar to what we observed in the *zim-1* mutant and the number was further increased upon removal of BRC-1-BRD-1 (*brc-1(xoe4); zim-2(tm574)*=4.91±0.96, *brd-1(ok1623); zim-2(tm574)*=4.86±0.93, *brc-1(xoe4); zim-3(xoe15)*=4.09±1.12, *brd-1(ok1623); zim-3(xoe15)*=3.83±1.10) (Fig 5A). Thus, BRC-1-BRD-1 limits the number of crossover precursors in spermatogenesis under circumstances where asynapsed chromosomes are present.

Previous analyses in oogenesis had indicated that when crossover formation is completely blocked by mutation of central components of the SC, COSA-1 accumulates at foci that represent aberrant recombination sites (Li *et al.* 2018; Woglar and Villeneuve 2018; Cahoon *et al.* 2019; Hurlock *et al.* 2020). We next examined GFP::COSA-1 in *syp-1* mutant males, in which germ cells fail to undergo chromosome synapsis and therefore do not form any interhomolog crossovers (Macqueen *et al.* 2002). As observed in hermaphrodites, *syp-1* mutant males exhibited a significant number of COSA-1 foci (4.0±1.20) (Fig. 5A). However, in the absence of BRC-1 or BRD-1, fewer GFP::COSA-1 foci were observed (*brc-1(xoe4); syp-1(me17)=*3.27±1.15, *brd-1(ok1623); syp-1(me17)=*3.56±1.51). This suggests that unlike the situation where crossover formation is inhibited on only a subset of chromosomes, BRC-1-BRD-1 promotes the localization of COSA-1 at recombination sites when no interhomolog crossovers are allowed to form.

### BRC-1 influences the genetic crossover landscape

Given the influence of BRC-1-BRD-1 on COSA-1 foci in the different mutants, we monitored genetic linkage between SNP markers on chromosomes *I* and *V* in male Bristol/Hawaiian hybrid strains to assess whether BRC-1-BRD-1 alters the formation of *bona fide* crossovers (Fig 6A). Inactivation of BRC-1 had little effect on the genetic map length of either chromosome *I* or *V* (*I*: WT=45.74cM; *brc-1(xoe4)*=52.17cM; *V*: WT=45.21cM; *brc-1(xoe4)*=50.82cM; Sup Table 4, Fig 6B). In *C. elegans*, crossovers are not evenly distributed along the length of the chromosomes but are enriched on the gene-poor arms (Barnes *et al.* 1995; Lim *et al.* 2008; Rockman and Kruglyak 2009). Similar to what we reported for oocytes (Li *et al.* 2018), there is a statistically significant alteration in the distribution of crossovers in the *brc-1* mutant on both chromosomes *I* and *V* compared to wild-type males (Sup Table 4, Fig 6C). In the *brc-1* mutant we observed an expansion in the center of the chromosome, with more crossovers in the center-right interval on chromosome *I* (30.21% vs.13.95%; p=0.0123) and the left-center interval on chromosome *V* compared to wild type (12.9% vs. 3.53%; p=0.0304) (Sup Table 4, Fig. 6C).

**Figure 6.**
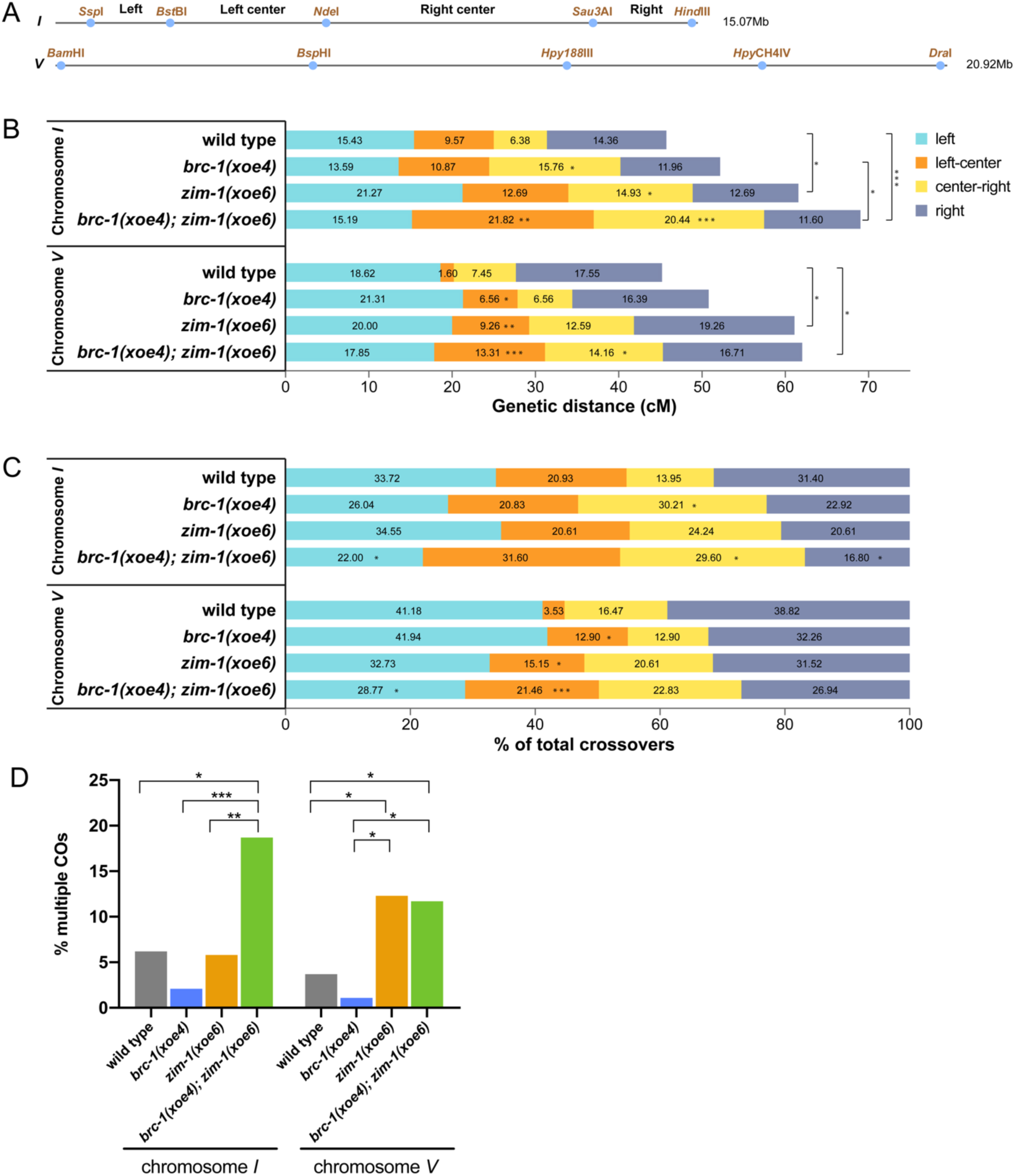
BRC-1 inhibits extra crossovers on chromosome *I* in the *zim-1* mutant. A) SNP markers on chromosome *I* and *V* used for genotyping; primers and additional information are included in Sup Table 2. B) CO frequency on chromosome *I* in wild type (n = 188), *brc-1(xoe4)* (n = 184), *zim-1(xoe6)* (n = 268) and *brc-1(xoe4); zim-1(xoe6)* (n = 362) mutants and on chromosome *V* in wild type (n = 188), *brc-1(xoe4)* (n = 183), *zim-1(xoe6)* (n = 270) and *brc-1(xoe4); zim-1(xoe6)* (n = 353) mutants. n = number of individuals analyzed per genotype. C) CO distribution among recombinants on chromosome *I* and *V* in wild type, *brc-1(xoe4), zim-1(xoe6)* and *brc-1(xoe4); zim-1(xoe6)* mutants. D) Percent of recombinant chromosomes containing multiple COs calculated as 100 x (DCO + TCOs)/(SCO + DCOs + TCOs). Statistical analyses were conducted using Fisher exact test on 2-by-2 contingency tables, * p<0.05; ** p<0.001; *** p<0.0001.

We next monitored linkage between SNP markers in the *zim-1* and *brc-1; zim-1* mutant males. We observed a significant increase in the genetic map length on both chromosome *I* and *V* in *zim-1* and *brc-1; zim-1* compared to wild-type males (*I*: *zim-1(xoe6)*=61.57cM p=0.0014, *brc-1(xoe4); zim-1(xoe6)*=69.06cM p=0.0001; *V*: *zim-1(xoe6)*=61.11cM p=0.0089, *brc-1(xoe4); zim-1(xoe6)*=62.04cM p=0.0024; Sup Table 4, Fig 6B). In addition to the expanded genetic maps, crossover distributions were also altered in these two mutants. The percentage of crossovers on the left and right arms of chromosome *I* were reduced in *brc-1; zim-1* compared to wild type (left: 22% vs. 33.7% p=0.0426; right: 16.8% vs. 31.4% p=0.0053), while the right-center interval was expanded in *brc-1; zim-1* compared to wild-type males (29.6% vs. 13.9% p=0.004; Sup Table 4, Fig 6C). On chromosome *V* there was an increased percentage of crossovers in the left-center interval in *zim-1* compared to wild-type males (15.2% vs. 3.5% p=0.0053), and it was further expanded in *brc-1; zim-1* (21.5% p=0.0001), while the right-center had significantly more crossovers in *brc-1; zim-1* compared to *brc-1* males (22.83% vs. 12.9% p=0.045; Sup Table 4, Fig 6C).

A unique feature of *C. elegans* oogenic meiosis is that on average there is a single crossover per chromosome pair per meiosis (Albertson *et al.* 1997; Hillers and Villeneuve 2003; Hammarlund *et al.* 2005). This is attributed to very strong interference, which is the phenomenon that the presence of one crossover at one position decreases the probability of formation of another crossover nearby. Analyses in spermatocytes also suggested that there is usually a single crossover per chromosome pair (Meneely *et al.* 2002; Kaur and Rockman 2014); however, Lim *et al.* (Lim *et al.* 2008) reported that interference was not as strong in male meiosis due to the appearance of closely spaced double crossovers (DCOs). We detected 5 DCOs on chromosome *I* and 3 DCOs on chromosome *V* in a total of 188 wild-type spermatocytes, which corresponds to 6.2% and 3.7% of total crossover events (Sup Table 4, Fig 6D). Fewer DCOs were detected in the *brc-1* mutant males, although this was not statistically different (chromosome *I*: 2 DCO/184, 2.1%; chromosome *V*: 1 DCO/183, 1.1%; Sup Table 4, Fig. 6D). In contrast, we previously detected no DCOs in 187 oocytes in either wild type or *brc-1* oocytes (Li *et al.* 2018). In the *zim-1* mutant, we detected 9 DCOs in 268 spermatocytes on chromosome *I*, which corresponds to 5.8% of total crossover events and is not significantly different compared to wild type; however, in the *brc-1; zim-1* double mutant, a significantly higher percentage of COs were DCOs and triple crossovers (TCOs): 37 DCOs and 2 TCOs were detected in 362 spermatocytes, which collectively is 18.7% of total CO events (Sup Table 4, Fig 6D). On chromosome *V, zim-1* had elevated levels of DCOs and TCOs (18/270, 12.3%) compared to wild type and *brc-1* spermatocytes, but this was not further increased in the *brc-1; zim-1* double mutant (23/353, 11.7%; Sup Table 4, Fig 6D).

Given the increased frequency of DCOs, we calculated interference. While most intervals had absolute interference of 1 in wild type and *brc-1*, the detection of DCOs resulted in decreased interference in two intervals on both chromosome *I* and chromosome *V* (Table 1). *zim-1* mutant males displayed reduced interference in all intervals except the left to left center and left center to right center intervals on chromosome *I*. Inactivation of BRC-1 in the *zim-1* mutant further impaired interference in all intervals on chromosome *I*, but had a variable effect on chromosome *V*, although they did not reach statistical significance (Table 1). Taken together, the elevated number of COSA-1 foci and increased numbers of DCOs and TCOs in the *brc-1; zim-1* mutant on chromosome *I* suggest that BRC-1-BRD-1 inhibits supernumerary crossovers on other chromosomes when asynapsed chromosomes are present.

**Table 1:** Crossover interference on chromosome *I* and *V*.

| WT (I) | exp DCO freq | obs DCO freq | c.o.c | interference |
| --- | --- | --- | --- | --- |
| L - LC | 0.0148 | 0.0160 | 1.0805 | -0.0805 |
| L - CR | 0.0098 | 0.0000 | 0.0000 | 1.0000 |
| L - R | 0.0222 | 0.0106 | 0.4802 | 0.5198 |
| LC - R | 0.0138 | 0.0000 | 0.0000 | 1.0000 |
| CR - R | 0.0092 | 0.0000 | 0.0000 | 1.0000 |
| LC - CR | 0.0061 | 0.0000 | 0.0000 | 1.0000 |
| WT (V) | exp DCO freq | obs DCO freq | c.o.c | interference |
| L - LC | 0.0030 | 0.0000 | 0.0000 | 1.0000 |
| L - CR | 0.0139 | 0.0000 | 0.0000 | 1.0000 |
| L - R | 0.0327 | 0.0106 | 0.3255 | 0.6745 |
| LC - R | 0.0028 | 0.0000 | 0.0000 | 1.0000 |
| CR - R | 0.0131 | 0.0053 | 0.4069 | 0.5931 |
| LC - CR | 0.0012 | 0.0000 | 0.0000 | 1.0000 |
| *brc-1* (I) | exp DCO freq | obs DCO freq | c.o.c | interference |
| L - LC | 0.0148 | 0.0000 | 0.0000 | 1.0000 |
| L - CR | 0.0214 | 0.0054 | 0.2538 | 0.7462 |
| L - R | 0.0162 | 0.0054 | 0.3345 | 0.6655 |
| LC - R | 0.0130 | 0.0000 | 0.0000 | 1.0000 |
| CR - R | 0.0188 | 0.0000 | 0.0000 | 1.0000 |
| LC - CR | 0.0171 | 0.0000 | 0.0000 | 1.0000 |
| *brc-1* (V) | exp DCO freq | obs DCO freq | c.o.c | interference |
| L - LC | 0.0140 | 0.0000 | 0.0000 | 1.0000 |
| L - CR | 0.0140 | 0.0000 | 0.0000 | 1.0000 |
| L - R | 0.0349 | 0.0055 | 0.1564 | 0.8436 |
| LC - R | 0.0107 | 0.0000 | 0.0000 | 1.0000 |
| CR - R | 0.0107 | 0.0000 | 0.0000 | 1.0000 |
| LC - CR | 0.0043 | 0.0000 | 0.0000 | 1.0000 |
| *zim-1* (I) | exp DCO freq | obs DCO freq | c.o.c | interference |
| L - LC | 0.0270 | 0.0000 | 0.0000 | 1.0000 |
| L - CR | 0.0317 | 0.0149 | 0.4702 | 0.5298 |
| L - R | 0.0270 | 0.0112 | 0.4149 | 0.5851 |
| LC - R | 0.0161 | 0.0037 | 0.2318 | 0.7682 |
| CR - R | 0.0189 | 0.0037 | 0.1971 | 0.8029 |
| LC - CR | 0.0189 | 0.0000 | 0.0000 | 1.0000 |
| *zim-1* (V) | exp DCO freq | obs DCO freq | c.o.c | interference |
| L - LC | 0.0185 | 0.0037 | 0.2000 | 0.8000 |
| L - CR | 0.0252 | 0.0148 | 0.5882 | 0.4118 |
| L - R | 0.0385 | 0.0333 | 0.8654 | 0.1346 |
| LC - R | 0.0178 | 0.0111 | 0.6231 | 0.3769 |
| CR - R | 0.0243 | 0.0037 | 0.1527 | 0.8473 |
| LC - CR | 0.0117 | 0.0074 | 0.6353 | 0.3647 |
| *brc-1; zim-1* (I) | exp DCO freq | obs DCO freq | c.o.c | interference |
| L - LC | 0.0332 | 0.0138 | 0.4166 | 0.5834 |
| L - CR | 0.0311 | 0.0304 | 0.9784 | 0.0216 |
| L - R | 0.0176 | 0.0110 | 0.6268 | 0.3732 |
| LC - R | 0.0253 | 0.0193 | 0.7637 | 0.2363 |
| CR - R | 0.0237 | 0.0110 | 0.4659 | 0.5341 |
| LC - CR | 0.0446 | 0.0331 | 0.7431 | 0.2569 |
| *brc-1; zim-1* (V) | exp DCO freq | obs DCO freq | c.o.c | interference |
| L - LC | 0.0238 | 0.0170 | 0.7153 | 0.2847 |
| L - CR | 0.0253 | 0.0085 | 0.3362 | 0.6638 |
| L - R | 0.0298 | 0.0227 | 0.7598 | 0.2402 |
| LC - R | 0.0223 | 0.0057 | 0.2546 | 0.7454 |
| CR - R | 0.0237 | 0.0085 | 0.3590 | 0.6410 |
| LC - CR | 0.0189 | 0.0028 | 0.1502 | 0.8498 |

L = left interval; LC = left-center interval; CR = center-right interval; R = right interval.
DCO: double crossover; expected DCO: (crossover frequency at interval "A") x
(crossover frequency at interval "B"). c.o.c. (coefficient of coincidence) = actual DCO
frequency/ expected DCO frequency; Interference = 1- c.o.c. See Sup Table 4 for data
used for calculations.

## Discussion

We show here that the BRC-1-BRD-1 complex functions in early processing of meiotic DSBs to promote HR and also inhibits supernumerary crossovers when some chromosomes are unable to form crossovers in male meiosis. These functions are distinct from previous analyses in oogenesis and suggests that this complex is differently regulated during male and female meiosis to optimize sperm versus oocyte production.

### Overlapping but distinct meiotic silencing pathways in *C. elegans* and mammals

Mouse BRCA1 is essential for MSCI, recruiting ATR for H2AX phosphorylation and chromosome compaction (Turner *et al.* 2004). ATR, in turn, promotes the accumulation of additional BRCA1 and other DNA damage signaling proteins to hemizygous regions of sex chromosomes, perhaps in response to unrepaired meiotic DSBs (Royo *et al.* 2013; Lu and Yu 2015). Accumulation of DNA damage response components are linked to the recruitment of SETDB1 methyltransferase for H3K9me3 enrichment and gene silencing (Hirota *et al.* 2018). While *C. elegans* ATR ortholog and to a lesser extent the related ATM checkpoint kinases are critical for targeting H3K9me2 to the hemizygous X chromosome in male germ cells, removal of BRC-1-BRD-1 had no effect on either the deposition of H3K9me2 or on gene silencing of the X chromosome (Fig 1), suggesting that BRC-1-BRD-1 does not mediate MSCI in *C. elegans* male meiosis.

As master regulators, ATR and ATM phosphorylate a large number of substrates (Matsuoka *et al.* 2007; Mu *et al.* 2007); consequently the observed effect on meiotic silencing may be indirect. Indeed, a recent study revealed that these kinases function in multiple aspects of meiotic recombination during *C. elegans* oogenesis (Li and Yanowitz 2019). We have shown that the X chromosome in males is refractory to ATM-dependent meiotic DSB formation feedback mechanisms (Checchi *et al.* 2014), suggesting that the defect in accumulation of H3K9me2 may not be through unrepaired DSBs, as is proposed in mammals. In addition, ATR is normally present at very low levels in the male germ line and is not enriched on the X chromosome, implying an indirect role for this kinase in MSCI, although it accumulates genome-wide in response to exogenous DNA damage or in mutants impaired for recombination or synapsis (Jaramillo-Lambert *et al.* 2010). Further, a *C. elegans* H2AX ortholog has not been identified that can be phosphorylated by ATR/ATM (Boulton 2006). On the other hand, the SETDB1 methyltransferase, MET-2, mediates H3K9me2 deposition and gene silencing of the X chromosome in male germ cells (Bessler *et al.* 2010; Checchi and Engebrecht 2011), analogous to SETDB1 function in mammals (Hirota *et al.* 2018). However, in contrast to mice, MET-2 does not accumulate on the X chromosome of male germ cells (Yang *et al.* 2019). Thus, the mechanisms whereby ATL-1/ATM-1 promote accumulation of H3K9me2 via MET-2 on the X chromosome of males remains to be elucidated but perhaps is linked to a small RNA pathway that is required for meiotic silencing (She *et al.* 2009). Nonetheless, the overlapping but distinct requirements for components that mediate MSCI in worms and mammals suggest that meiotic silencing is a conserved feature of meiosis in metazoans; however, the pathways used to target repressive chromatin marks have evolved independently.

### BRC-1-BRD-1 regulates DSB processing to promote HR in male germ cells

In somatic cells, BRCA1-BARD1 functions in DNA damage signaling and repair to promote genome integrity (Kouznetsova *et al.* 2009; Li and Greenberg 2012; Savage and Harkin 2015; Takaoka and Miki 2018). Critical to the maintenance of the genome is the choice of pathways for repair of DSBs: HR, NHEJ and other pathways including microhomology mediated end joining. Whether HR or NHEJ is used is largely driven by DNA end resection. Several studies support the hypothesis that BRCA1-BARD1 regulates the choice between repair by HR and NHEJ. Initial evidence for this was based on the observation that *brca1*^*-/-*^ embryonic lethality can be rescued by removal of 53BP1, a DNA damage response protein that promotes NHEJ (Cao *et al.* 2009; Bouwman *et al.* 2010; Bunting *et al.* 2010). More recent work has suggested that BRCA1-BARD1 promotes DNA end resection by removing a chromatin barrier through ubiquitination of histone H2A (Densham *et al.* 2016) and/or through speeding up resection by interaction with CtIP, a protein that promotes end resection (Cruz-Garcia *et al.* 2014). Studies by other groups also showed that BRCA1 and CtIP work together with the MRX/N complex to mediate resection of complex breaks, including Spo11-dependent meiotic DSBs (Hartsuiker *et al.* 2009; Aparicio *et al.* 2016).

Our analyses of male meiosis reveal that similar to the role of BRCA1-BARD1 in somatic cells, this complex regulates the processing of meiotic DSB to promote repair by HR in male meiosis. First, in the absence of BRC-1-BRD-1, fewer RAD-51 foci were observed in meiotic prophase (Fig 2), suggesting BRC-1-BRD-1 functions at or prior to RAD-51 loading onto resected ends. Further we show that the reduction in RAD-51 foci can be suppressed by mutation of NHEJ proteins, consistent with a role of BRC-1-BRD-1 in regulating the choice between HR and NHEJ. Further, the localization of BRC-1 to foci in early meiotic prophase, which presumably represent sites of ongoing recombination, is dependent on DNA resection (Fig 3). These findings point to a role for BRC-1-BRD-1 in promoting repair by HR, likely by regulating resection. It is important to note that *brc-1* and *brd-1* mutants exhibit only subtle meiotic phenotypes, in contrast to the phenotypic consequence of removing components of the resection machinery. Mutation of CtIP (*C. elegans* COM-1) or components of the MRX/N complex leads to high embryonic lethality and almost a complete absence of RAD-51 loading (Chin and Villeneuve 2001; Hayashi *et al.* 2007; Lemmens *et al.* 2013; Girard *et al.* 2018). Thus, while BRC-1-BRD-1 is not essential for resection, our data is consistent with this complex regulating resection speed or extent, as in somatic cells.

### BRC-1-BRD-1 function when male meiosis is perturbed

In addition to a role in processing of meiotic DSBs to promote HR, we show that BRC-1-BRD-1 also functions to promote progeny viability when male meiosis is perturbed under conditions when some chromosome pairs fail to pair, synapse and form a crossover (Fig 4). While this is also true for female meiosis, the phenotypic consequences of inactivating BRC-1 or BRD-1 are distinct in the sexes. During female meiosis, removal of BRC-1-BRD-1 under checkpoint activating conditions leads to premature disassembly of the RAD-51 filament (Li *et al.* 2018); however, no effect on RAD-51 filament stability was detected during male meiosis, despite observing fewer RAD-51 foci presumably due to the role of BRC-1-BRD-1 in DSB end processing as is the case in an otherwise wild-type background (Fig 4). Additionally, while the crossover landscape is altered in both male and female meiosis, opposite effects of removing BRC-1-BRD-1 in the *zim-1* mutant were observed. In female meiosis mutation of *brc-1* or *brd-1* in the *zim-1* background led to fewer COSA-1-marked crossover designation events, while during male meiosis the numbers increased. One possibility to explain this observation is that destabilization of the RAD-51 filament in mid-late pachytene in female meiosis leads to fewer meiotic recombination intermediates that can be processed into COSA-1-marked crossover precursors.

Analysis of genetic crossovers on chromosome *I* and *V* in male meiosis revealed an altered crossover distribution in the *brc-1* mutant, such that more crossovers occurred at the chromosome center, and fewer on the arms, as was previously observed in oogenesis (Li *et al.* 2018). Alteration in crossover distribution in the *brc-1* mutant may result from changes in the chromatin landscape, which has been linked to BRCA1 function in mammals (Broering *et al.* 2014; Densham *et al.* 2016), and has been shown to alter crossover patterning (Mezard *et al.* 2015; Yu *et al.* 2016). A surprising number of *C. elegans* meiotic mutants display altered crossover distribution (Zetka and Rose 1995; Meneely *et al.* 2012; Saito *et al.* 2012; Saito *et al.* 2013; Wagner *et al.* 2010; Chung *et al.* 2015; Hong *et al.* 2016; Jagut *et al.* 2016; Janisiw *et al.* 2020). While the underlying mechanisms are not clear, it may be a consequence of altering the use of crossover versus non-crossover pathways within a particular chromatin environment as suggested by Saito and Colaiacovo (Saito and Colaiacovo 2017).

Under conditions where not all chromosomes can form a crossover (e.g., *zim-1*), we observed differential effects in the presence and absence of BRC-1 on chromosomes *I* versus *V*. On chromosome *I*, the *zim-1* mutant had elevated SCOs, but not DCOs compared to wild type, while removal of BRC-1 in the *zim-1* mutant resulted in elevated levels of DCOs at the expense of SCOs. We propose that this reflects a shift from three and four strand DCOs, which are included in the SCO class and are not marked by COSA-1, in *zim-1*, to two-strand DCOs marked by COSA-1 in *brc-1; zim-1*. In contrast, on chromosome *V*, the *zim-1* mutant showed significantly higher levels of DCOs compared to wild type but removing BRC-1 had little effect. During female meiosis, inactivation of BRC-1 in the *zim-1* mutant background had the opposite effect, in which decreasing numbers of DCOs and elevated numbers of SCOs were observed on chromosome *V*, presumably due to a shift from two strand DCOs to three and four strand DCOs (Li *et al.* 2018). Thus, there are both chromosome-specific and sex-specific effects on the crossover landscape when BRC-1 is inactivated. The chromosome-specific effect may be a consequence of size; chromosome *I* is the smallest chromosome, while chromosome *V* is the largest chromosome. Recent work in yeast suggests that small chromosomes use multiple mechanisms to ensure the formation of the obligate crossover (Murakami *et al*. 2020). Therefore, the differential impact on chromosome *I* versus *V* may be due to the mechanisms in place to promote crossover formation on small chromosomes. Alternatively, other chromosome-specific features may influence which DSBs are converted into crossovers when BRC-1-BRD-1 is not present to constrain extra crossover formation during male meiosis.

Why does removal of BRC-1-BRD-1 enhance embryonic lethality when a subset of chromosomes fails to form a crossover? Mutation of *brc-1* enhanced crossover distribution defects as well as the number of DCOs on some chromosomes in the *zim-1*mutant background (Fig 6). Alteration in crossover position (Altendorfer *et al.* 2020) as well as elevated crossover numbers (Hollis *et al.* 2020) are deleterious during *C. elegans* meiosis. This is likely a consequence of the holocentric nature of *C. elegans* chromosomes and the requirement to establish asymmetric domains as defined by the single crossover site for accurate cohesion release and chromosome segregation (De Carvalho *et al.* 2008; Ferrandiz *et al.* 2018). Additionally, BRC-1 has an established role in promoting inter-sister HR (Adamo *et al.* 2008; Garcia-Muse *et al.* 2019). In the absence of BRC-1-BRD-1, resected breaks on chromosomes that cannot undergo crossover formation may fail to be repaired prior to the meiotic divisions, leading to chromosome fragmentation, loss of genetic material and aneuploid gametes.

### Sex-specific regulation of meiosis

Our analyses of BRC-1-BRD-1 reveals several differences between male and female meiosis. First, while there is currently no direct measure of DSB formation in *C. elegans*, we detected a higher average of RAD-51 foci in male versus female germ cells, suggesting that more DSBs are induced in spermatocytes. Usage of DSBs hotspots in mice has also revealed sex-specific differences (Brick *et al.* 2018). Second, BRC-1-BRD-1 functions at different steps of meiotic recombination in the sexes. In males, BRC-1-BRD-1 influences the early processing of DSBs to promote HR, while in females, BRC-1-BRD-1 is engaged in mid-late pachytene to promote repair of breaks processed and assembled with RAD-51 by inter-sister recombination (Adamo *et al.* 2008). How BRC-1-BRD-1 is differentially regulated in the sexes is not known, but the spatiotemporal pattern of BRC-1-BRD-1 function mirrors MAP kinase activation in the male (transition zone/early pachytene) and female (mid-late pachytene) germ lines (Lee *et al.* 2007). Thus, MAP kinase and/or other signaling pathways could regulate the complex in a sex-specific manner to drive ubiquitination of different substrates in spermatogenesis versus oogenesis.

Overall *C. elegans* male meiosis appears to be less tightly regulated compared to female meiosis. For example, we detected a few DCOs in wild-type male meiosis, but none in oocytes (Li *et al.* 2018). Further, previous analyses have shown that males lack germ line apoptosis, one mechanism to enhance gamete quality by removing defective or damaged germ cells (Gartner *et al.* 2000; Jaramillo-Lambert *et al.* 2010). Interestingly, the same checkpoint proteins that promote apoptosis in females in response to meiotic errors, function to promote meiotic fidelity in males (Jaramillo-Lambert *et al.* 2010). Why male meiosis appears to lack regulatory mechanisms yet has a reduced frequency of meiotic errors compared to oogenesis is currently unknown. Future analyses of *C. elegans* male meiosis may provide insight into the mechanisms that contribute to the fidelity of male gametes.

## Acknowledgements

We thank the Caenorhabditis Genetic Center, which is funded by NIH Office of Research Infrastructure Programs (P40 OD010440) for providing strains. We are grateful to Ben Mallory for generating *zim-3(xoe15)*, Jonathan Amezquita for generating *zim-1(xoe6)* and Lauren Ahmann, Arshdeep Kaur and Tara Shahrvini for help with construction of strains. We thank Takamune Saito and Marina Martinez-Garcia for advice on meiotic mapping experiments and the Engebrecht lab for thoughtful discussions. This work was supported by National Institutes of Health GM103860 and GM103860S1 to JE.

## Figure Legends

**Supplemental Figure 1. Analysis of RAD-51 foci in *brc-1(tm1145) brd-1(dw1), brd-1(ok1623), cku-70(tm1524)* and double and triple mutant male germ lines.** A) Quantification of RAD-51 in indicated regions of the germ line. Box whisker plots show number of RAD-51 foci per nucleus in the different regions. Horizontal line of each box represents the median, top and bottom of each box represents the median of upper and lower quartiles, lines extending above and below boxes indicate standard deviation and individual data points are outliers from 5–95%. Statistical comparisons by Mann-Whitney of WT versus *brc-1(tm1145) brd-1(dw1), brc-1(tm1145) brd-1(dw1)* versus *brc-1(tm1145) brd-1(dw1)cku-70(tm1524)* and *brd-1(ok1623)* versus *brd-1(ok1623) cku-70(tm1524)* in the different regions of the germ line; *** p<0.0001, ** p<0.001, * p<0.05. All statistical comparisons are shown in Sup Table 3. PZ =proliferative zone; TZ = transition zone; EP = early pachytene; MP = mid-pachytene; LP = late pachytene. Number of nuclei scored in each region from a minimum of 3 germ lines: WT: PZ = 1225; TZ = 449; EP = 296; MP = 251; LP = 226; *brc-1(tm1145) brd-1(dw1)*: PZ = 869; TZ = 628; EP = 435; MP = 328; LP = 336; *brc-1(tm1145) brd-1(dw1)cku-70(tm1524)*: PZ = 634; TZ = 361; EP = 329; MP = 329; LP = 289; *cku-70(tm1524)*: PZ = 869; TZ = 382; EP = 403; MP = 292; LP = 323; *brd-1(ok1623) cku-70(tm1524)*: PZ = 418; TZ = 188; EP = 124; MP = 124; LP = 117; *brd-1(ok1623)*: PZ = 758; TZ = 349; EP = 358; MP = 269; LP = 301; B) Representative images of nuclei from indicated genotypes at late pachytene of the germ line stained with antibodies against RAD-51 (red) and counterstained with DAPI. Scale bar=5μm

**Supplemental Figure 2. Early meiotic GFP::BRC-1 foci partially co-localize with RAD-51.** A) Representative images of transition zone/early pachytene region of the germ line fixed and stained with antibodies against GFP (green) to detect GFP::BRC-1 and RAD-51 (red), counterstained with DAPI (blue). Arrows point to overlap between GFP::BRC-1 and RAD-51 foci. Scale bar = 10 μm. B) High-magnification images of live worms expressing GFP::BRC-1 in *mre-11(ok179)* null (PZ = proliferative zone, TZ = transition zone, EP = early pachytene, MP = mid-pachytene, LP = late pachytene, DP = diplotene). Images are projections through half of the gonad. Scale bar=5μm.

